# A Rationally Designed Transgene Drives CAR T Functional Persistence and Durable Regression of Solid Tumors

**DOI:** 10.64898/2026.09.18.752754

**Authors:** Rupesh H. Amin, Kevin G. Haworth, Howell F. Moffett, Jerry C. Chen, Lisa Park, Jason K. Yokoyama, Allan L. Wang, Leah J. Tait, Maria M. Steele, John T. Crowl, Thaddeus M. Davenport, Joseph DeSautelle, Jared L. Hammer, Willimark M. Obenza, Vanessa R. Montoya, Robin L. Kirkpatrick, Tina Tan, Kristie M. Shirley, Bradley Hammerson, Robert A. Langan, David S. Clausen, Paul J. Sample, Brian D. Weitzner, Shujun Yuan, Marc J. Lajoie, Scott E. Boyken, Aaron E. Foster

## Abstract

The eradication of solid tumors by chimeric antigen receptor (CAR) T cells requires dynamic therapies capable of outlasting an immune suppressive tumor microenvironment (TME). However, biological barriers—including rapid exhaustion, poor expansion, and loss of stem-like memory—quickly neutralize these therapies^1^. Because single-technology interventions often introduce unacceptable tradeoffs between efficacy and safety, durable remission demands a paradigm where multiple engineered solutions work in concert. To holistically address these mechanisms, we rationally designed a single-vector transgene that intrinsically drives CAR T functional persistence. The platform integrates four synergistic technologies: a high-avidity mesothelin (MSLN)-targeting CAR optimized to resist shed decoy antigens, a T-cell activation-responsive promoter (OUTLAST OP1) resisting exhaustion, a CD8α-targeted designed IL-2 cytokine (OUTSMART dIL-2) driving intratumoral CAR-T expansion, and an EGFRopt^2^ safety switch. In lung and ovarian cancer models, this rational integration was required to drive antigen-dependent T cell expansion and eradicate established tumors at extremely low CAR T doses. Furthermore, engineered cells established a self-renewing pool of stem-like memory T cells capable of rejecting tumor rechallenge months later. Ultimately, this work demonstrates that intrinsic T cell dysfunction and extrinsic tumor-derived barriers can be simultaneously overcome by integrating synergistic technologies.

## Introduction

Chimeric antigen receptor (CAR) T cell therapies have achieved remarkable clinical success in hematological malignancies^3,4^, but this success has not translated to solid tumors^5^ due to an immune suppressive tumor microenvironment (TME) that impairs T cell function and survival. Clinical evidence has established a suite of compounding mechanisms driving therapeutic failure: poor tumor infiltration^6,7^, progressive functional exhaustion and loss of T cell stemness due to chronic antigen exposure^8,9^, susceptibility to immune suppressive TME signaling^1^, and the neutralization of engineered receptors via shed decoy antigens^10–13^. Initial efforts to counteract these barriers have focused on optimizations that boost potency, such as increasing CAR activity/expression or incorporating constitutively-expressed cytokines to provide pro-survival signals^14^. While these strategies have achieved some benefit, they often rely on static or constitutive signaling that introduces new liabilities, such as accelerated exhaustion or systemic cytokine-mediated toxicities^15^. These limitations underscore that overcoming solid tumor barriers requires moving beyond static, single-component modifications toward engineering approaches that capitalize on the T cell’s inherent capacity for dynamic, context-dependent decision making. We hypothesized that fully unleashing this potential against solid tumors would require a multi-component transgene, with each element intentionally designed to sense and conditionally overcome a specific failure mechanism at the tumor site.

Here, we describe the design and validation of a functionally persistent CAR T cell therapy for mesothelin (MSLN)-expressing cancers. MSLN is a glycoprotein that is highly-expressed on the surface of numerous solid tumors with high unmet need, including ovarian cancers, pancreatic cancers, and mesotheliomas^16^. Our approach combines four distinct technologies (**Figure 1**): i) a high-avidity, immune synapse-optimized MSLN CAR; ii) a T-cell activation-responsive promoter (OP1) to regulate transgene expression and mitigate T cell exhaustion; iii) a CD8α-targeted, designed IL-2 cytokine to drive antigen-dependent expansion of effector cells at the tumor; and iv) an optimized EGFR-based safety switch (EGFRopt^2^) that enables antibody-based T cell elimination. We demonstrate that this integrated system achieves complete and durable tumor regression at ultra-low CAR T doses in preclinical animal models, establishing new strategies for more completely addressing the challenges facing solid tumor efficacy.

**Figure 1.**
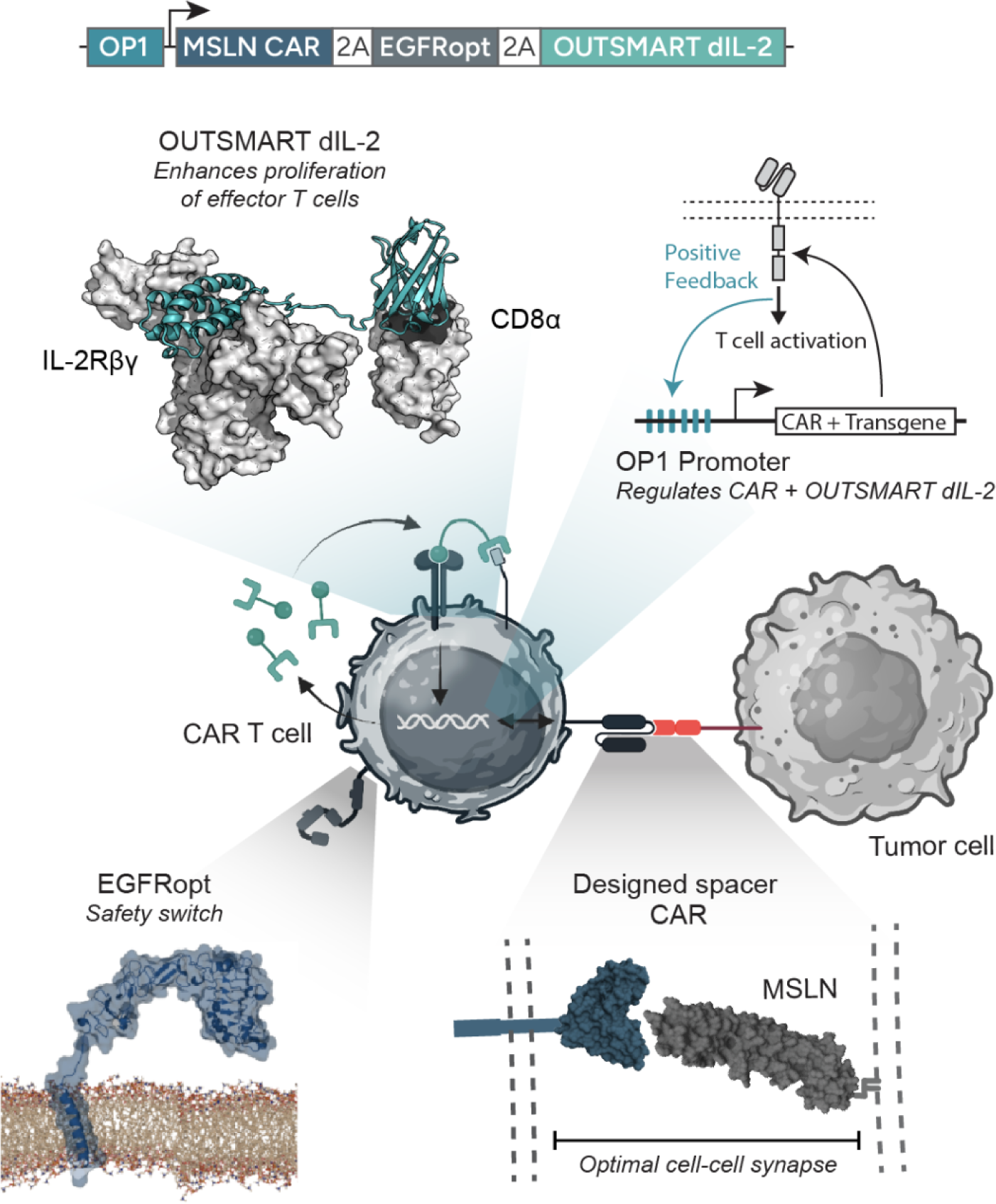
Integration of four distinct technologies that work in concert to overcome challenges to solid tumor efficacy: A high-avidity mesothelin (MSLN)-targeting CAR, which includes an optimized cell-cell synapse via tuning the spacer domain; a T-cell activation-inducible promoter (OUTLAST OP1) that resists exhaustion, as well as controls inducible expression of the other transgenes; a CD8α-targeted designed cytokine, OUTSMART designed IL-2 (dIL-2) that drives intratumoral CAR-T expansion; and an optimized safety switch (EGFRopt). Illustrations of cells were created using BioRender (https://biorender.com, Fontana, T. 2024).

### Engineering a high-avidity CAR T cell for potent tumor recognition

We first set out to create an optimal MSLN CAR with improved anti-tumor activity and resistance to inhibition due to soluble MSLN, which is shed from tumor cells. CAR T cell activity is partly dependent on the distance and geometry of the cell-cell synapse between the effector T cell and tumor cell^17^. Matching spacer length to target epitope location (longer for membrane-proximal, shorter for membrane-distal) is required to optimize immunological synapse distance and maximize CAR potency. To address this, we designed a set of CAR spacer sequences comprising a wide range of lengths (43.2 - 248.4 Å), derived from natural-occurring human immune proteins with proline and cysteine motifs that impart structural rigidity (**Figure 2a**). The MSLN extracellular domain (ECD) has an extended conformation comprising three subdomains (Regions I-III, where Region III is closest to the membrane and Region I is most distal) extending ∼90 Å away from the membrane (**Extended Data Fig. 1a**). To evaluate the interplay between epitope location and spacer geometry, we paired a panel of MSLN-targeting scFv and VHH binders (directed against Region I or III of the MSLN ECD) combinatorially with our engineered spacers, constructing CARs with a CD28 transmembrane domain and a 4-1BB/CD3ζ signaling endodomain. CAR constructs were screened in a Nur77-Jurkat reporter line to exclude binders driving antigen-independent tonic signaling, and subsequently evaluated in primary human T cells for target-dependent killing and cytokine production using an *in vitro* H1650 lung adenocarcinoma co-culture assay. CARs targeting the membrane-distal Region I of MSLN exhibited higher overall potency than those targeting juxtamembrane regions (**Extended Data Fig. 1b**). As expected for a distal binding epitope, Region I specific CARs were most effective when paired with shorter spacers, such as osp1 (43.2 Å), osp6 (64.8 Å), and osp8 (75.6 Å) (**Extended Data Fig. 1b,c**). Notably, the Region I-binding SS1 CAR, which was tested previously in human clinical trials, was the most potent amongst those screened but only when paired with a shorter osp1 spacer, which enhanced its performance over the CD8α spacer that was used clinically (**Extended Data Fig. 1c**). Additional fine-tuning of osp1 by subtracting Gly-Ser linker amino acids further improved performance of the SS1 CAR (**Extended Data Fig. 1d)**. Collectively, these data demonstrate the importance of the spacer domain and immune synapse as a critical component of CAR-T cell potency.

**Figure 2.**
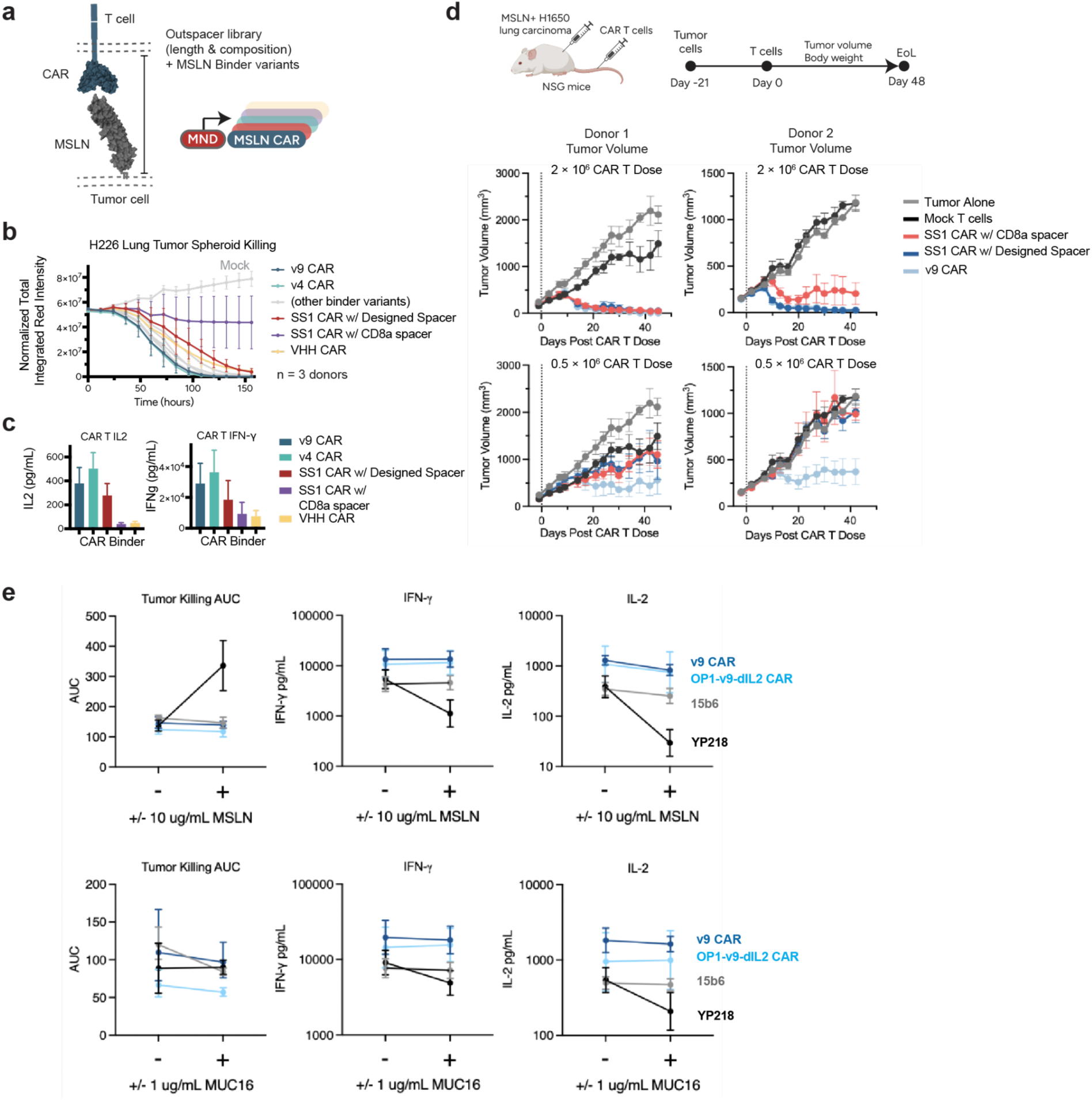
MSLN CAR binder and spacer optimization results in superior in vitro and in vivo efficacy. **a,** Schematic of CAR spacer design approach. **b**, The v9 CAR shows superior killing of H226 lung tumor spheroids over time. **c**, v9 CAR T cells produce high levels of IL-2 and IFN-γ upon tumor recognition. **d**, In a subcutaneous H1650 lung cancer model, the v9 CAR demonstrates improved tumor control compared to a benchmark CAR at a dose of 0.5 × 10^6^ cells. **e**, v9 CAR and full OP1-v9-dIL2 are resistant to soluble MSLN and MUC16; CAR 15b6 binds the juxtamembrane domain of MSLN and has been previously shown to be resistant to soluble MSLN; CAR YP218 shows inferior tumor killing and cytokine production in the presence of soluble MSLN or MUC16.

To further enhance potency, top binder/spacer leads underwent humanization and were screened in an *in vitro* H226 MSLN^+^ lung carcinoma spheroid co-culture assay (Methods). Humanized SS1 scFv variants (v4 and v9) formatted with the osp1 spacer demonstrated superior tumor spheroid killing (**Figure 2b**) and elevated secretion of IFN-γ and IL-2 (**Figure 2c**) relative to the original SS1 CAR (with CD8α or osp1 spacers) and the lead VHH construct (VHH-8). Next, CAR constructs were evaluated in immune-deficient NSG mice engrafted with MSLN^+^ H1650 tumors and treated with mock T or MSLN CAR-T cells. All CAR-T cells were effective at controlling tumors at 2 × 10^6^ cells/animal, but the v9 CAR was superior at lower cell doses (0.5 × 10^6^ cells/animal) (**Figure 2d**).

Previous work on MSLN-targeting therapeutics has highlighted soluble MSLN and MUC16 (the native ligand of MSLN) shed from tumor cells as potential barriers to efficacy due to their ability to act as competitors for CAR binding^10–13^. Others have attempted to mitigate this issue by engineering CARs that specifically target the juxtamembrane domain of MSLN left behind after shedding or RIII of MSLN that is not involved in MUC16 binding^10,11,18^. Compared to the parental SS1 scFv, the v9 scFv—which conferred superior CAR T cell performance *in vivo*—exhibited lower binding affinity for MSLN, driven primarily by a faster off-rate (k_off_) (**Extended Data Fig. 2**). We hypothesized that a faster dissociation rate would reduce receptor neutralization by shed antigen (soluble MSLN or MUC16). To test this, CAR T cell cytotoxicity and cytokine production were evaluated in co-culture assays with MSLN^+^ NCI-H1650 lung adenocarcinoma and H2052 mesothelioma cell lines—selected specifically for their lack of endogenous MSLN or MUC16 shedding—supplemented with recombinant soluble MSLN (10 μg/mL) or MUC16 (1 μg/mL). The v9 CAR maintained full cytotoxic and secretomic potency in the presence of either soluble competitor (**Figure 2e**). CAR T cells bearing the juxtamembrane-binding 15b6 scFv were unaffected by shed antigen, but secreted significantly lower levels of IL-2 and IFN-γ than v9 CAR T cells. In contrast, a previously published Region III-binding CAR (YP218^19^) showed impaired activity in the presence of soluble MSLN and MUC16, despite targeting an epitope distinct from the MUC16 binding site (**Figure 2e**). Collectively, these results demonstrate that optimizing binder kinetic properties and spacer geometry can overcome the inhibitory effects of shed decoy antigens while targeting the membrane-distal Region I of MSLN.

### OUTLAST OP1 T-cell activation-inducible promoter confers resistance to exhaustion

T cell exhaustion is an intrinsic state of dysfunction in which chronic CD3 signaling induces epigenetic changes that lock in a hypofunctional phenotype^20^. Previous studies have shown that transient “rest” from chronic antigen stimulation through the addition of the tyrosine kinase inhibitor dasatinib^21^, or by placing the CAR under endogenous regulatory elements of the TCR via knock-in at the TRAC locus^22^, can delay or mitigate CAR T cell exhaustion^21^. We hypothesized that dynamically regulating CAR expression using an antigen-responsive promoter could prevent exhaustion by mitigating chronic receptor signaling, without requiring exogenous small molecules or site-specific gene editing.

We generated a T cell activation responsive promoter, named OUTLAST Promoter 1 (OP1), that can regulate CAR expression in response to antigen-driven signaling. The OP1 promoter is composed of 10 repeats of consensus binding motifs for NFkB followed by a minimal promoter and 5’ untranslated region from human alpha-globin. The NF-kB signaling pathway is strongly activated by both the 4-1BB and CD3ζ signaling domains present in the CAR receptor^23,24^. The OP1 promoter drives increased transgene expression upon T cell activation, creating a positive feedback loop in response to antigen-dependent signaling (**Figure 3a-b**), and reduced expression in the absence of antigen. Importantly, while CAR T cells driven by a conventional constitutive promoter (MND) suffered a severe loss of function upon chronic antigen exposure, OP1-driven CAR T cells maintained robust cytokine secretion following repeated rounds of stimulation and mounted a significantly stronger response to acute challenge(**Figure 3c**). This functional benefit was consistent with a less-exhausted phenotype, and OP1-driven CAR T cells showed substantially lower expression of the inhibitory receptors PD-1 and TIGIT (**Figure 3d)**. The resistance to T cell exhaustion conferred by the OP1 promoter also improved tumor control in the MSLN^+^ H1650 xenograft mouse model (**Figure 3e**). To evaluate whether the OP1-driven functional benefit is generalizable to other CARs and tumor models, we tested it using a different MSLN CAR with a nanobody (VHH)-based binder against H1650 xenografts, as well as with a ROR1 CAR against a ROR1-expressing H1975 non-small cell lung adenocarcinoma xenograft^25^. In all three cases, OP1-regulated CAR T cells outperformed MND-expressed CAR T cells, demonstrating greater *in vivo* tumor control, peripheral blood expansion, and intratumoral expansion (**Extended Data Fig. 3**).

**Figure 3.**
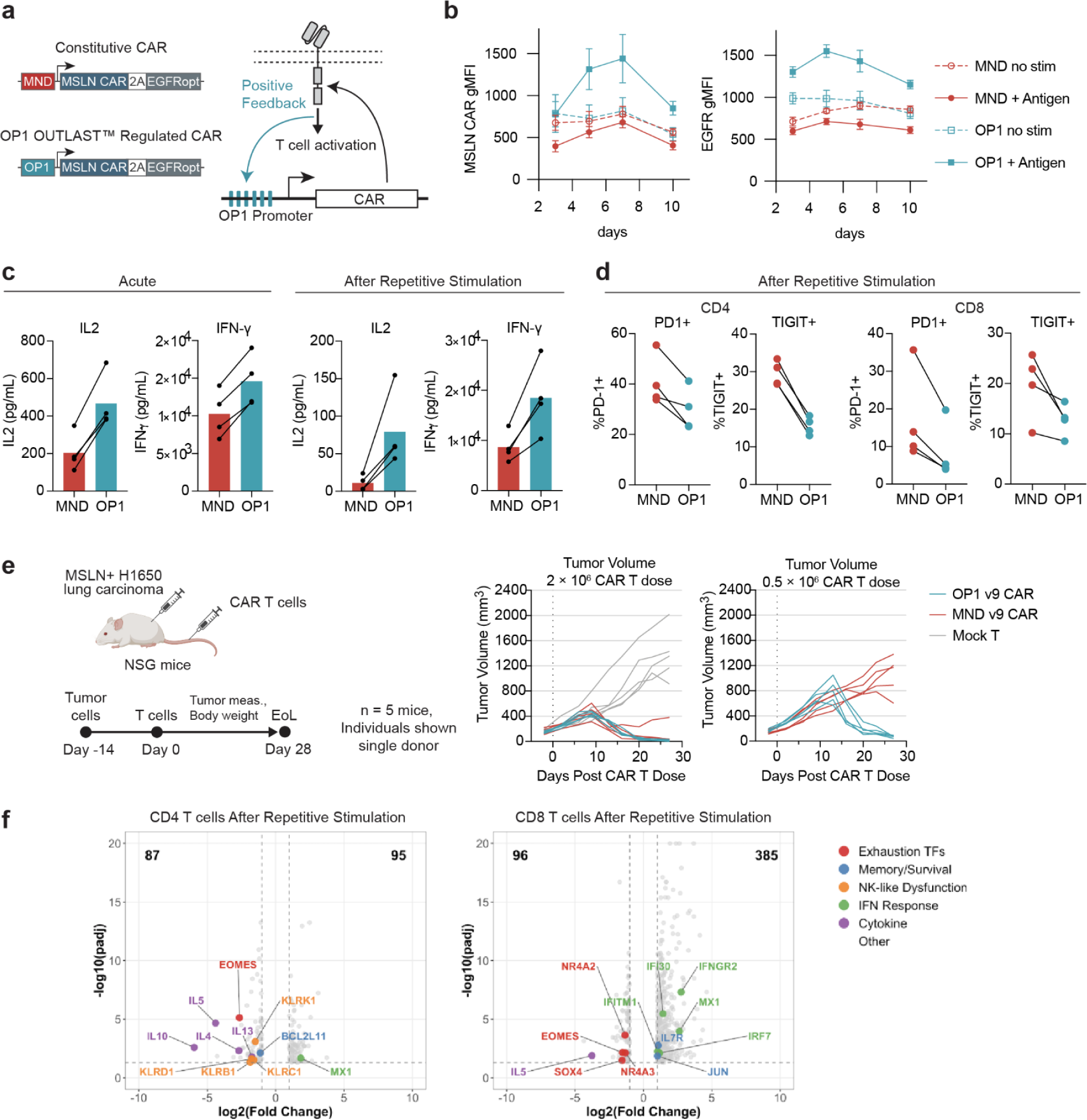
The OUTLAST OP1 promoter confers resistance to antigen-driven exhaustion. **a**, schematic of OP1 promoter positive feedback activity and constructs used to evaluate OP1. **b**, CAR expression levels over time during repetitive antigen stimulation or no stimulation. **c,** OP1-driven CAR T cells maintain cytokine production at higher levels after acute and repetitive stimulation compared to MND-driven CAR T cells. **d**, OP1-driven CAR T cells exhibit lower PD-1 and TIGIT expression after repetitive stimulation compared to MND-driven cells. **e**, In the H1650 model, OP1-driven CAR T cells show improved efficacy at a dose of 0.5 × 10^6^ cells compared to MND-driven CAR T cells. **f,** RNA-seq analysis shows OP1 CAR T cells have a less exhausted transcriptomic signature after repetitive stimulation; positive log2(fold-change) indicates genes higher in OP1 CAR, negative log2(fold-change) indicates genes higher in MND CAR.

To determine how the OP1 promoter preserves CAR T cell function, we analyzed the transcriptomes of OP1- and MND-regulated VHH-8 CAR T cells under baseline, acute, and chronic antigen challenge conditions by RNA sequencing (Methods). Repetitively stimulated OP1 CAR T cells resisted the NR4A/EOMES-driven exhaustion program, retained memory-associated IL-7R and JUN expression, maintained interferon response gene expression, and avoided the NK-like dysfunction phenotype (KLR gene upregulation) observed in repetitively stimulated MND CAR T cells (**Figure 3f, Extended Data Fig. 4)**.While dynamic regulation of CAR expression enhances T cell fitness by allowing the T cells to rest, positive feedback through the promoter’s NF-κB regulatory elements likely also contributes to these functional benefits. Sustained NF-κB signaling enforces a protective transcriptional program by upregulating BACH2 (which antagonizes cJun/BATF to prevent terminal differentiation)^26^, IL7R/CD127 (which supports memory stem cell formation and STAT5-mediated survival)^27^, and TOX2/BATF3 (which maintains stemness while repressing exhaustion drivers such as MAF, EOMES, and NR4A2/3^28,29^. Additionally, the 4-1BB costimulatory domain is known to activate noncanonical NF-κB signaling via the TRAF2/5-NIK-IKK pathway^23^, and CD3ζ signaling activates canonical NF-κB through PKC-mediated IKK phosphorylation^24^. Further studies will be required to fully elucidate these mechanisms, but the functional benefit of OP1 is clear, demonstrated by enhanced CAR T cell performance across multiple CARs and solid tumor models in vitro and in vivo.

### OUTSMART dIL-2 cytokine drives robust, antigen-dependent expansion

Having established the anti-exhaustion transcriptional profile and modularity of the OP1 promoter using the VHH-8 CAR, we next sought to construct our final, all-in-one optimized candidate. For this integrated circuit, we selected the v9 scFv MSLN CAR, which had demonstrated the highest overall potency and resistance to shed antigen in our earlier binder/spacer optimization screens (**Figure 2**). We therefore integrated OUTSMART designed IL-2 (dIL-2)[manuscript in preparation]—a CD8α-targeted designed cytokine—into the OP1-driven circuit alongside the v9 CAR and the EGFRopt safety switch (**Figure 4a**). This all-in-one design makes production of OUTSMART dIL-2 antigen-dependent, concentrating it at tumor sites, and increasing CAR T cell expansion and persistence, while maintaining the stemness and counter-exhaustion benefits of OP1 CAR regulation. In vitro, these “OP1-v9-dIL2” CAR T cells demonstrated antigen responsive production of dIL-2 cytokine across repeated cycles of stimulation and rest (**Figure 4b**). We next evaluated the OP1-v9-dIL2 construct in NSG mouse xenograft models. The absence of an endogenous immune system in these mice provides an isolated environment to explicitly demonstrate that the combination of OUTLAST OP1 and OUTSMART dIL-2 intrinsically drives CAR T expansion, preserves stemness, and establishes durable functional persistence. The efficacy of OP1-v9-dIL2 was compared to the v9 CAR alone driven by either MND or OP1 promoter in subcutaneous H1650 lung adenocarcinoma (**Figure 4c-d**) and intraperitoneal SKOV3 ovarian adenocarcinoma (**Figure 4e-f**). In both models, OP1-v9-dIL2 eradicated tumors at a cell dose of 0.5 × 10^6^ CAR T, a dose where the MND v9 CAR alone failed (**Figure 4c,e**). Compared to the OP1 v9 CAR (which lacks dIL-2), OP1-v9-dIL2 drove increased CAR T expansion and more consistent tumor control between donors and at lower cell doses (**Extended Data Fig. 5-6**). In both models, OP1-v9-dIL2 demonstrated efficacy down to a dose of 0.05 × 10^6^ CAR T. The superior efficacy and robust CAR T cell expansion in the peripheral blood was associated with preservation of naive/stem-like CD8+ CAR-T cells (CD45RA^+^CD62L^+^) suggestive of long-term functional persistence (**Figure 4d**). We assessed levels of OP1-regulated OUTSMART dIL-2 in peripheral blood (**Extended Data Fig. 5-6**) during CAR T mediated tumor regression. In H1650 xenograft experiments, levels of serum OUTSMART dIL-2 increased over time, reaching a peak of ∼200 pg/ml at the highest CAR T dose. The time of peak OUTSMART-dIL2 corresponded to the maximum tumor volume, after which the OUTSMART dIL-2 levels declined as tumor burden decreased (**Extended Data Fig. 5**). This same correlation between OUTSMART dIL-2 levels and tumor volume was also observed in the SKOV3 model (**Extended Data Fig. 6**).

**Figure 4.**
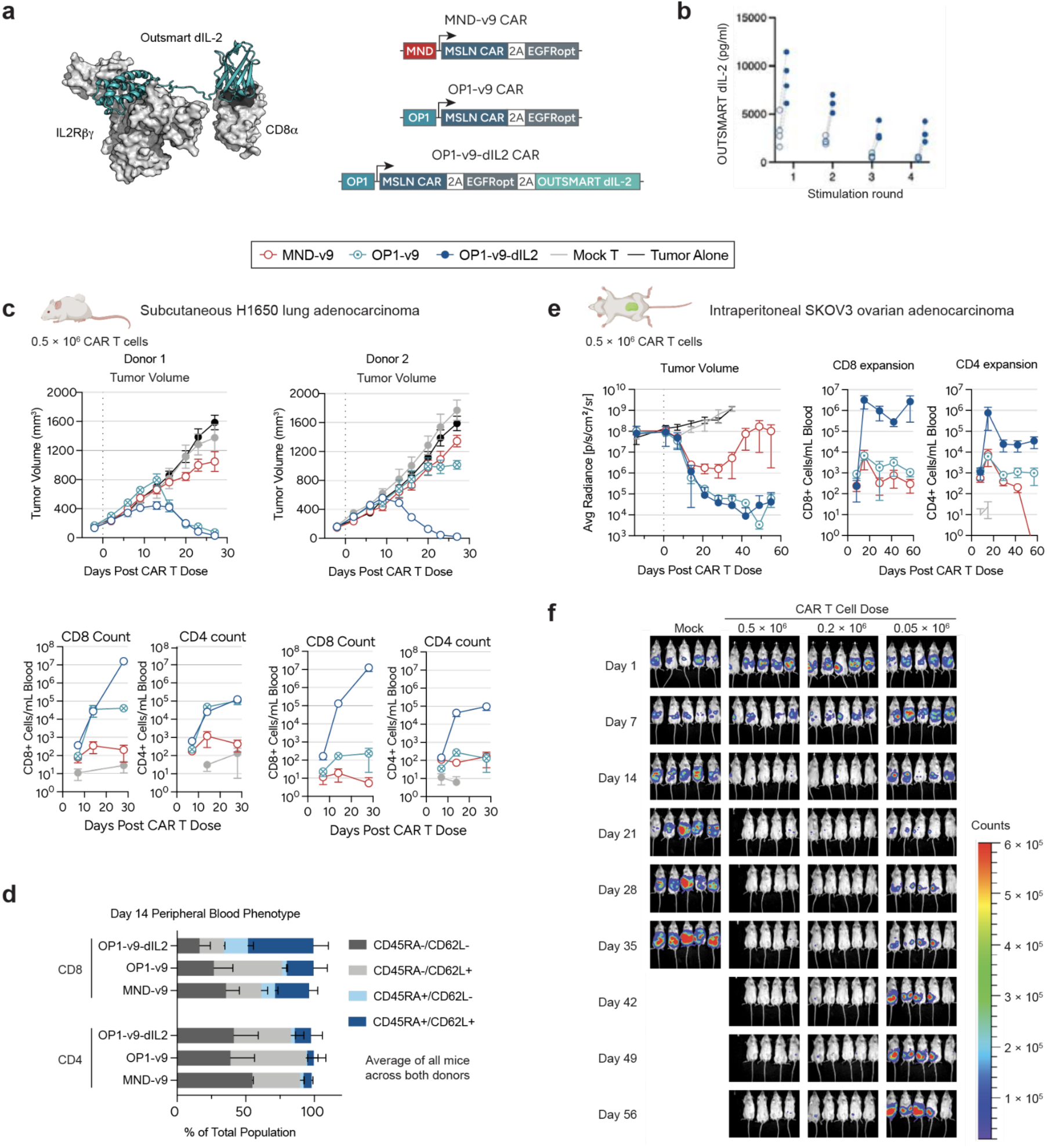
OP1-v9-dIL2 CAR T cells achieve complete tumor regression at very low cell doses. **a,** Schematic of the full OP1-v9-dIL2 construct, which integrates the OUTSMART dIL-2 designed cytokine. **b**, in vitro levels of inducible OUTSMART dIL-2 secretion (pg/ml) in response to repeat antigen stimulation; CAR T cells made from 4 healthy donors, each donor shown as an individual point, with the same donor connected by lines. In both subcutaneous H1650 lung (**c-d**) and intraperitoneal SKOV3 ovarian cancer models (**e-f**), OP1-v9-dIL2 drives massive T cell expansion and eradicates tumors at doses where therapies lacking any single component fail. **c**, In vivo tumor control and T cell counts from peripheral blood for OP1-v9-dIL2 at a cell dose of 0.5 × 10^6^ across two independent blood donors; representative of 4 donors tested (**Extended Data** Fig. 5). **d**, CAR T cell phenotype measured by flow cytometry, CD45RA/CD62L. **e,** In vivo tumor control and T cell counts from peripheral blood for SKOV3 ovarian cancer model; single representative donor shown out of 4 total donors tested (**Extended Data** Fig. 6). **f,** Imaging of individual mice from (e), using firefly luciferase (ffLuc) for direct in vivo imaging of tumors using a luminescent based detection instrument. Days listed are in reference to post-CAR T cell dosing.

### OUTLAST OP1 and OUTSMART dIL-2 in concert establish a self-renewing pool of stem-like memory T cells

To evaluate the long-term persistence and recall potential of OP1-v9-dIL2 CAR T cells, H1650 tumor-bearing mice were treated with low doses (0.2 × 10^6^ or 0.05 × 10^6^ CAR T cells). At both dose levels, primary tumors were completely cleared within 50 days, coinciding with robust peripheral CAR T cell expansion. To test recall responses, mice in the 0.05 × 10^6^ dose cohort were rechallenged on day 70 post-treatment with a secondary contralateral H1650 tumor injection (**Figure 5a**). Persisting CAR T cells rapidly rejected the secondary tumors (**Figure 5b**), exhibiting a pronounced re-expansion in peripheral blood relative to non-rechallenged control mice, in which CAR T cell counts continued to contract in the absence of antigen stimulation (**Figure 5c**). Phenotypically, OP1-v9-dIL2 CAR T cells maintained a high proportion of stem-like/naive (CD45RA+ CD62L+) cells throughout the study, while dynamically expanding effector memory and terminal effector subsets upon tumor rechallenge (**Figure 5d**).

**Figure 5.**
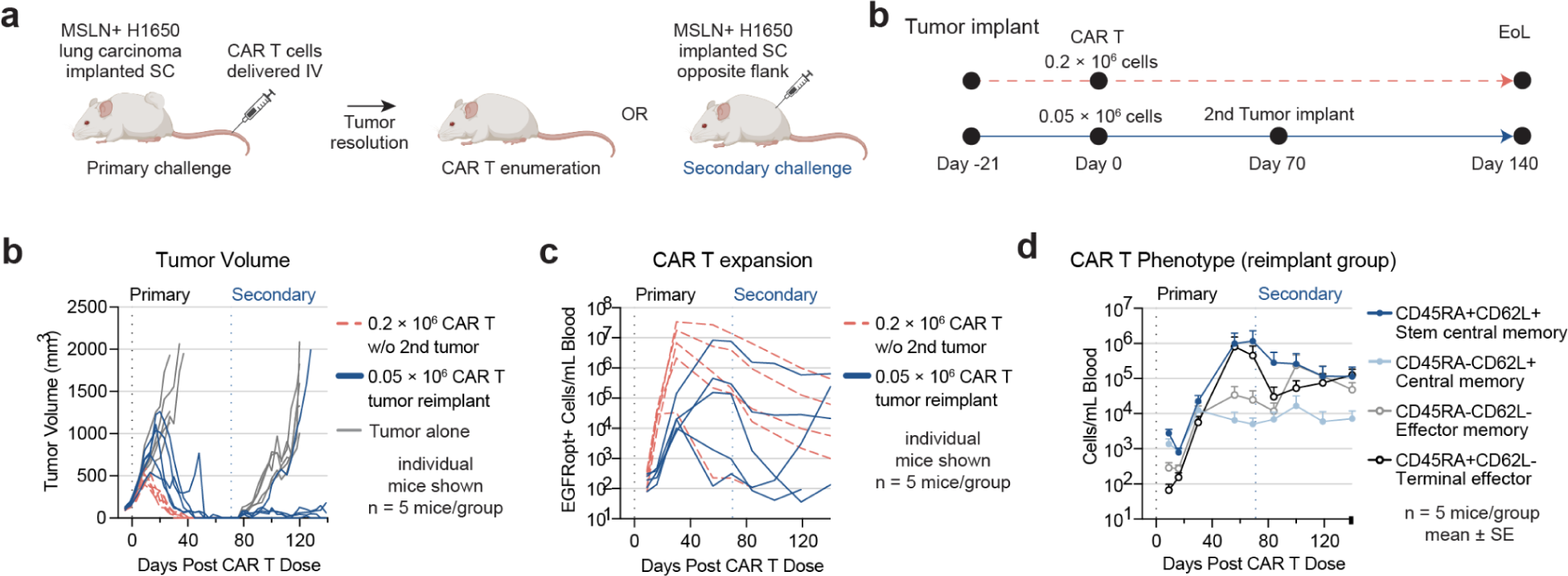
OP1-v9-dIL2 establishes a persistent, functional, and stem-like memory T cell pool. **a,** Mice that cleared a primary H1650 tumor with OP1-v9-dIL2 CAR T cells are protected from a secondary tumor challenge at day 70, associated with re-expansion of a Tscm-phenotype population. **b,** CD8+ T cells isolated from primary recipients after tumor clearance are functional and can eradicate tumors upon serial transfer to secondary recipients, and still maintain **c,** CAR T cell proliferative capacity and **d,** a more naive/stem-memory like phenotype.

To determine whether the persistent CD45RA+ CD62L+ CAR T cell population represents a functional reservoir of stem-like T cells capable of sustained self-renewal and recall over an extended period, we evaluated their performance under the high stringency of a serial transfer model. Primary mice bearing H1650 subcutaneous tumors were treated with 0.2 × 10^6^ CAR T cells; all mice cleared their primary tumors by day 40 (**Figure 6b**), accompanied by robust peripheral CD8+ CAR T cell expansion (**Figure 6c**). On day 40 post-treatment, CAR T cells were isolated from the peripheral blood of primary cured mice via cheek bleed and adoptively transferred at doses of 0.5 × 10^6^ or 0.2 × 10^6^ into secondary recipients bearing large, established H1650 tumors averaging 250 to 500 mm3 (**Figure 6a**). Upon transfer, these experienced CAR T cells robustly re-expanded, eliminating tumors in all secondary mice at the 0.5 × 10^6^ dose and in 3 of 4 mice at the 0.2 × 10^6^ dose (**Figure 6b,c**). Notably, OP1-v9-dIL2 CAR T cells preserved a high proportion of CD45RA+ CD62L+ cells in secondary recipients throughout the study (**Figure 6d**). Collectively, these data demonstrate that following tumor clearance, OP1-v9-dIL2 maintains a highly functional, stem-like CAR T cell pool capable of long-term self-renewal and vigorous expansion upon repeated exposure to heavy tumor burden.

**Figure 6.**
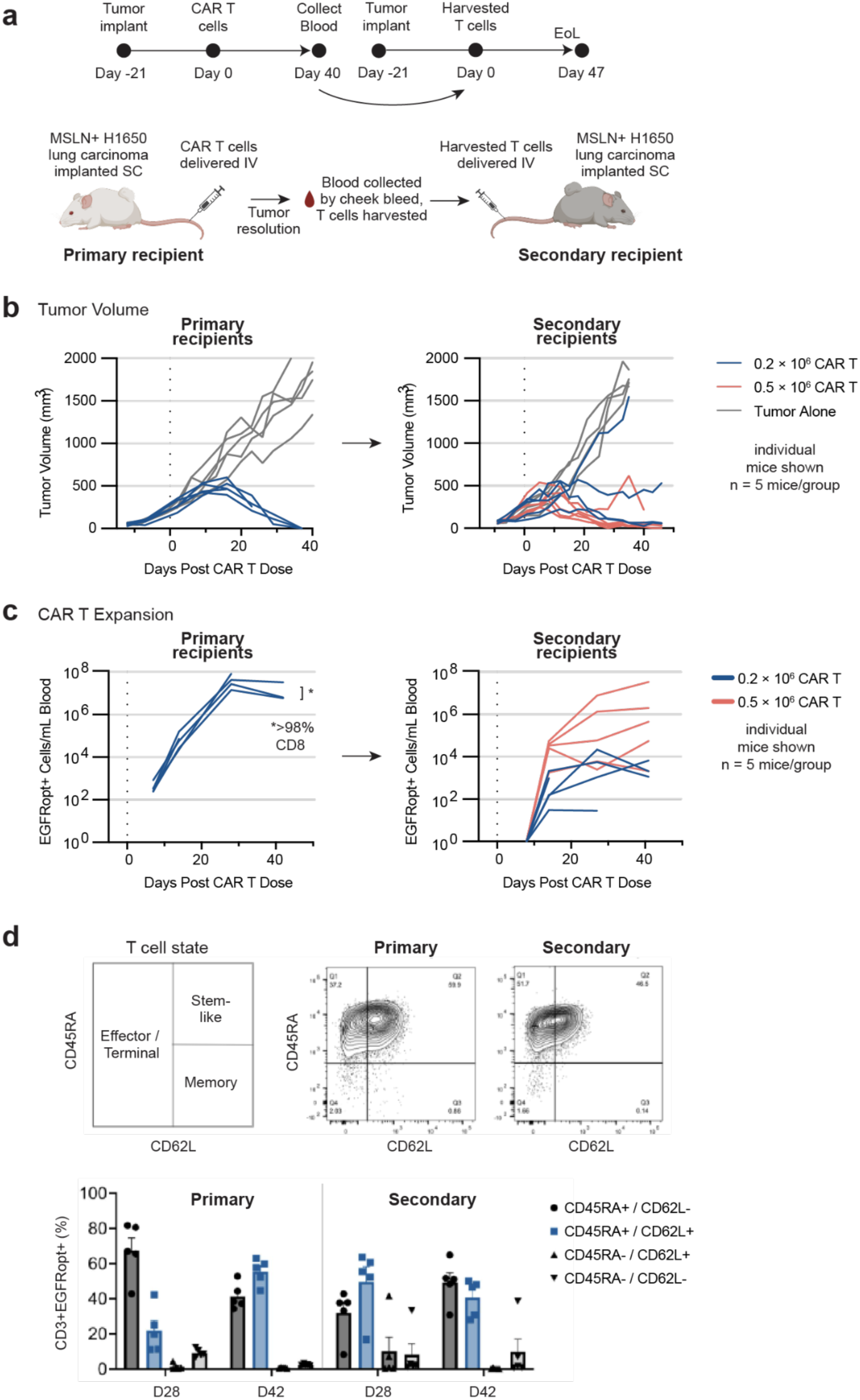
OP1-v9-dIL2 CD8 CAR T cells re-expand with antigen challenge without T cell help and maintain anti-tumor activity. **a,** NSG mice with SC H1650 tumors were administered 0.2 × 10^6^ CAR T cells IV; after tumor resolution, blood was harvested from primary mice and peripheral blood CAR T isolated for IV delivery to secondary tumor bearing recipients at 0.2 × 10^6^ and 0.5 × 10^6^ T cell doses (transferred T cells from primary mice were >98% CD8). **b,** tumor volume control. **c-d,** In the second set of mice, OP1 CAR T cells still maintain **c,** CAR T cell proliferative capacity and **d,** a more naive/stem-memory like phenotype.

### MSLN specificity and sensitivity

We evaluated the safety, specificity, and antigen sensitivity of the full platform. MND-v9 CAR T cells completely eliminated tumor lines expressing high to moderate levels of MSLN (including lung carcinoma lines NCI-H226 and NCI-H1650, and ovarian cancer lines OVCAR3 and SKOV3), but displayed only modest cytotoxic activity against low-expressing lines such as A549 lung carcinoma (**Extended Data Fig. 7a**). These CAR T cells failed to kill tumor cells expressing little to no MSLN (e.g., K562 and NCI-H1975), but regained full cytotoxicity when these lines were engineered to overexpress MSLN (**Extended Data Fig. 7b**). Incorporating the OP1 promoter and dIL2 module did not alter this specificity profile; OP1-v9-dIL2 CAR T cells exhibited killing activity comparable to MND-v9 across all tested lines (**Extended Data Fig. 7a,b**). Cytokine secretion (IL-2 and IFN-gamma) mirrored this antigen dependence, with high-MSLN lines triggering the strongest responses, low-to-moderate lines inducing reduced secretion, and MSLN-negative lines eliciting no cytokine release above mock-transduced controls (**Extended Data Fig. 7c**). Finally, antigen-dependent specificity was confirmed in vivo using a bilateral dual-tumor xenograft model. OP1-v9-dIL2 CAR T cells selectively cleared MSLN-positive H1650 subcutaneous tumors while sparing MSLN-low H1975 tumors on the contralateral flank of the same animals (**Extended Data Fig. 8**).

### Incorporation of EGFRopt safety switch

Finally, OP1-v9-dIL2 also includes an EGFRopt safety switch which can be used to eliminate the CAR T cells through administration of EGFR-targeting antibodies such as cetuximab^2^. Administration of cetuximab *in vivo* to OP1-v9-dIL2 CAR T cell-engrafted mice led to the rapid depletion of CAR T cells, providing a mechanism to control the therapy if needed. At the first time point evaluated 48 hours after cetuximab administration, an average 90% reduction in CAR T cell numbers was observed with all cetuximab dose levels. CAR T cell numbers continued to decrease further over the time course compared to mice administered CAR T cells without cetuximab (**Extended Data Fig. 9**).

## Discussion

Here we have detailed the design and preclinical validation of a CAR T cell therapy platform that synergistically combines four distinct technologies to address key barriers to durable efficacy in solid tumors. By integrating an optimized MSLN CAR with a regulated promoter that reduces exhaustion and a targeted cytokine that promotes expansion, OP1-v9-dIL2 achieves complete and durable tumor regression at very low cell doses in aggressive tumor models. Consistent with the functional persistence observed in rechallenge mouse xenograft experiments, OP1-v9-dIL2 establishes a long-lived pool of functional, stem-like memory T cells, a critical feature for preventing tumor relapse^27^.

Rather than relying on the optimization of a single component in isolation, we demonstrate the power of a modular, multi-component approach in which distinct technologies are layered to overcome specific biological barriers. The OP1 promoter mitigates the toxicity and exhaustion risks of constitutive signaling, while the designed cytokine provides sustained cytokine signaling required for robust expansion^30^, but does so in a localized, antigen-dependent manner that minimizes systemic exposure. Furthermore, while the immunodeficient xenograft models utilized in this study highlight the profound intrinsic benefits of this circuit, the fully secreted OUTSMART dIL-2 was intentionally designed to extend these effects to the broader tumor microenvironment. This CD8α-targeted cytokine preferentially stimulates bystander CD8+ T cells and Natural Killer (NK) cells while minimizing the activation of immunosuppressive regulatory T cells. Together, these findings indicate that the OP1-v9-dIL2 platform not only programs autonomous CAR T cell persistence, but also possesses the capacity to recruit and activate endogenous anti-tumor immunity. By combining protein design with synthetic biology, this platform holistically addresses clinically-validated mechanisms that limit solid tumor CAR T cell therapy, demonstrating that both intrinsic T cell dysfunction and tumor-derived barriers can be simultaneously defeated through the rational integration of synergistic technologies.

## Acknowledgements

We thank Eric Soller, Stan Riddell and Margo Roberts for helpful discussions and feedback on the manuscript. We thank current and former members of the Outpace Bio team, including Jen Running Deer, Stephanie Zimmerman and Lesley Jones for assistance with molecular cloning, Erik Hermans for assistance with cloning and lentivirus production, and Andrew Ng for helpful discussions during the early stages of this project.

## Author Contributions

R.A., A.E.F., M.J.L, S.E.B., K.G.H., and H.F.M. conceived of the work in this study. K.G.H. and J.C.C. designed and performed the *in vivo* studies. L.P. and J.Y. performed *in vitro* functional screening of CAR constructs and binder-spacer pairs. A.W., L.J.T., M.S., J.T.C., J.H., and R.K. performed in vitro functional assays and analyzed data. T.T. performed bulk cell preps and provided technical support. W.O. performed flow cytometry assays and provided technical support; K.S. performed molecular cloning and assisted with functional assays; B.H. performed protein production and reagent generation. T.M.D., S.Y., and B.H. analyzed binding kinetics data. T.M.D, J.D., and S.Y. generated and optimized VHH and scFv binders. B.D.W., R.A.L., and S.E.B. performed computational protein design for the dIL-2 cytokine component; P.J.S., J.T.C., R.A. and H.M. analyzed RNA-seq data; D.S.C. performed statistical analyses. V.R.M. and A.W. produced lentivirus. A.E.F., S.E.B., R.A., K.G.H., and H.M. wrote the paper, and all authors reviewed and edited the manuscript. A.E.F., M.J.L., and S.E.B. supervised the work.

## AI Use Disclosure Statement

Gemini (Google) was used to assist with grammar and sentence structure in this manuscript. The author(s) reviewed and edited the output and take full responsibility for the content of the published article.

## Extended Data Figures

**Extended Data Figure 1.**
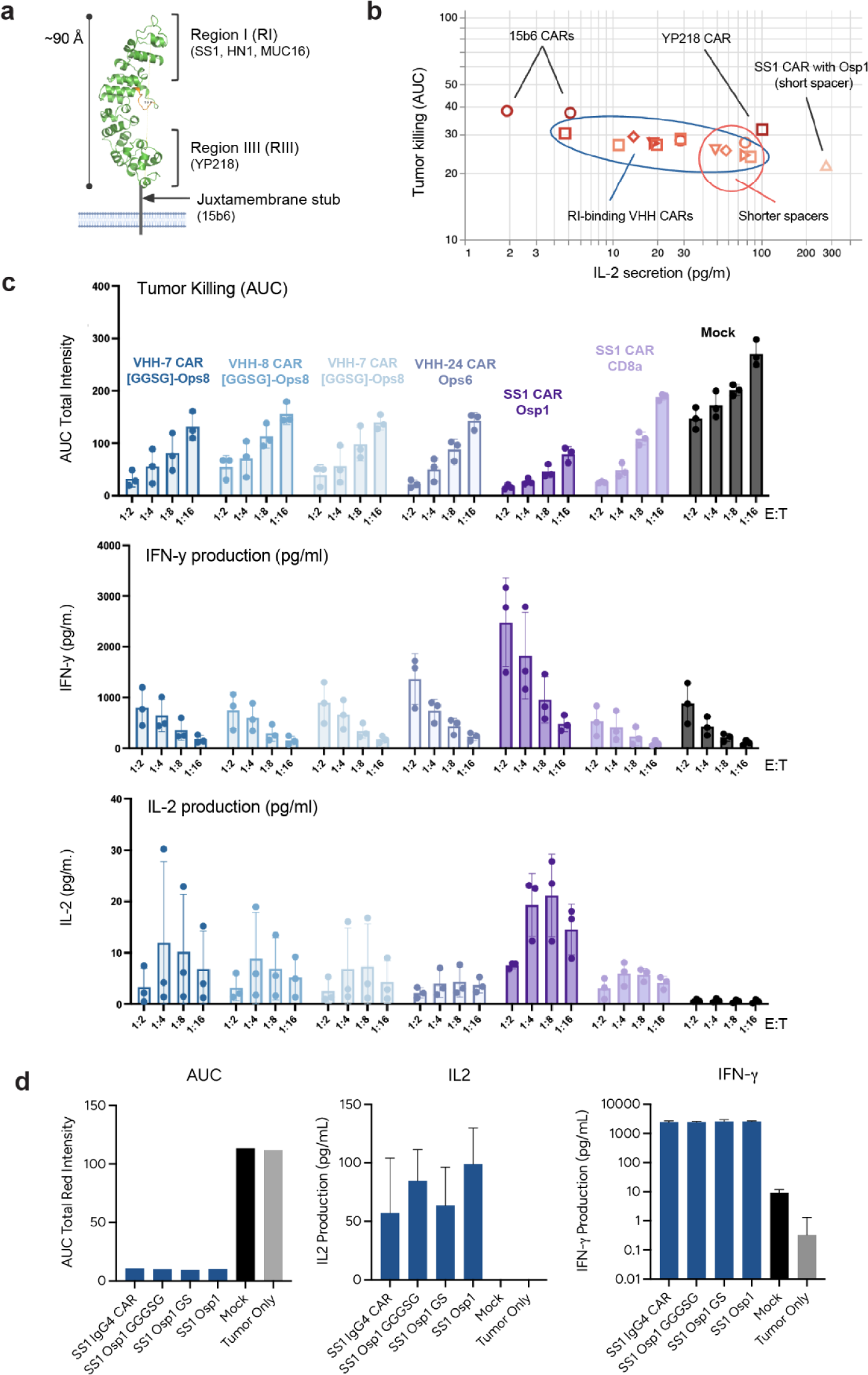
(Goes with Figure 2). Screening of CAR binder-spacer pairs. **a,** Structural model of the MSLN ECD. **b-d,** *in vitro* functional screens of CAR binder-spacer pairs measuring tumor cell killing and cytokine secretion in a tumor co-culture assay using a H1650 lung adenocarcinoma cell line. Tumor killing measured by Incucyte, area under the curve (AUC; lower value indicates better killing).

**Extended Data Figure 2.**
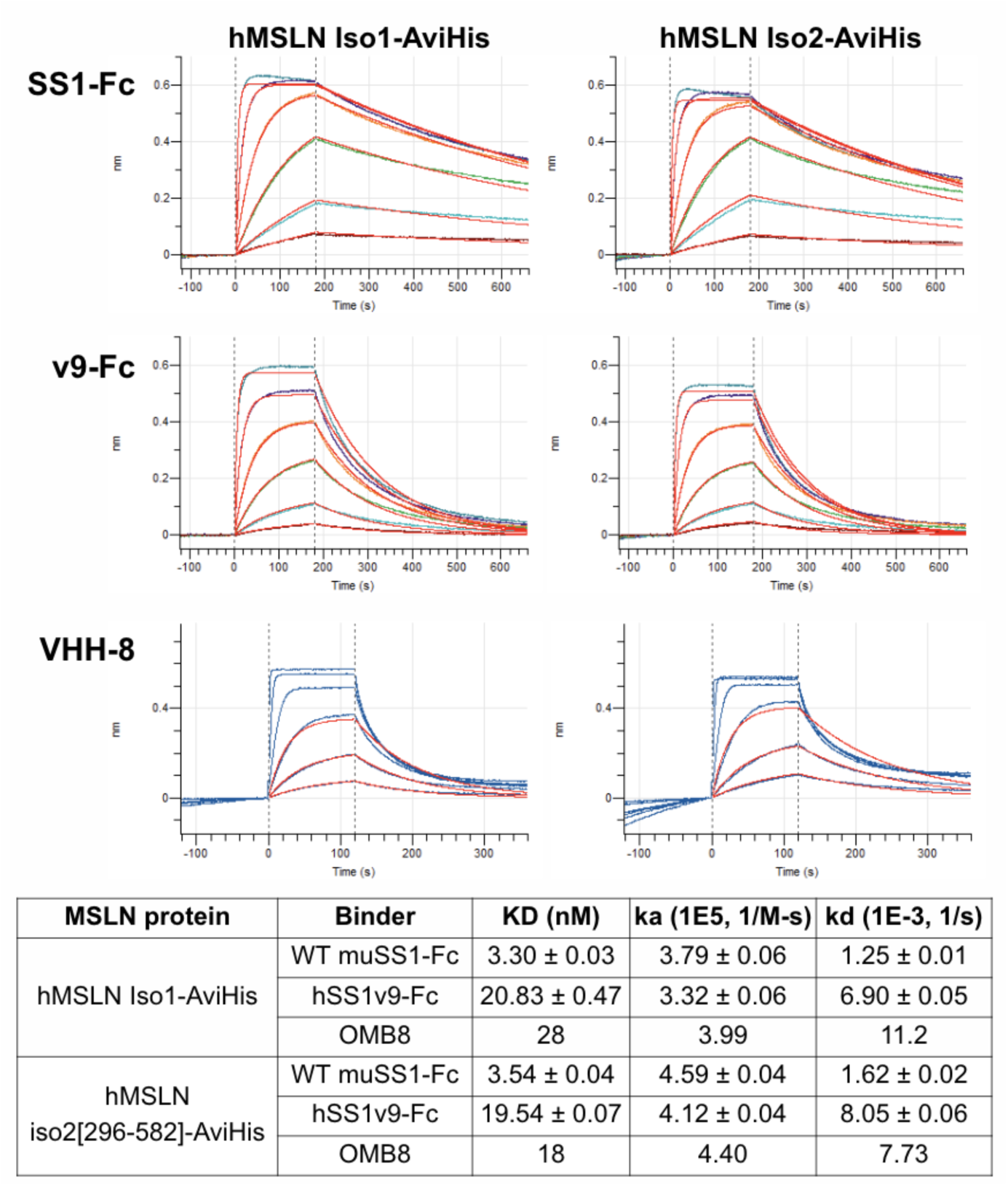
Binding kinetics measured by biolayer interferometry (BLI, Octet) for the MSLN binders used in the CAR constructs of this study, with human biotinylated AviHis-tagged MSLN ECD isoform 2 (“hMSLN Iso2-AviHis”) immobilized on Octet tips.

**Extended Data Figure 3.**
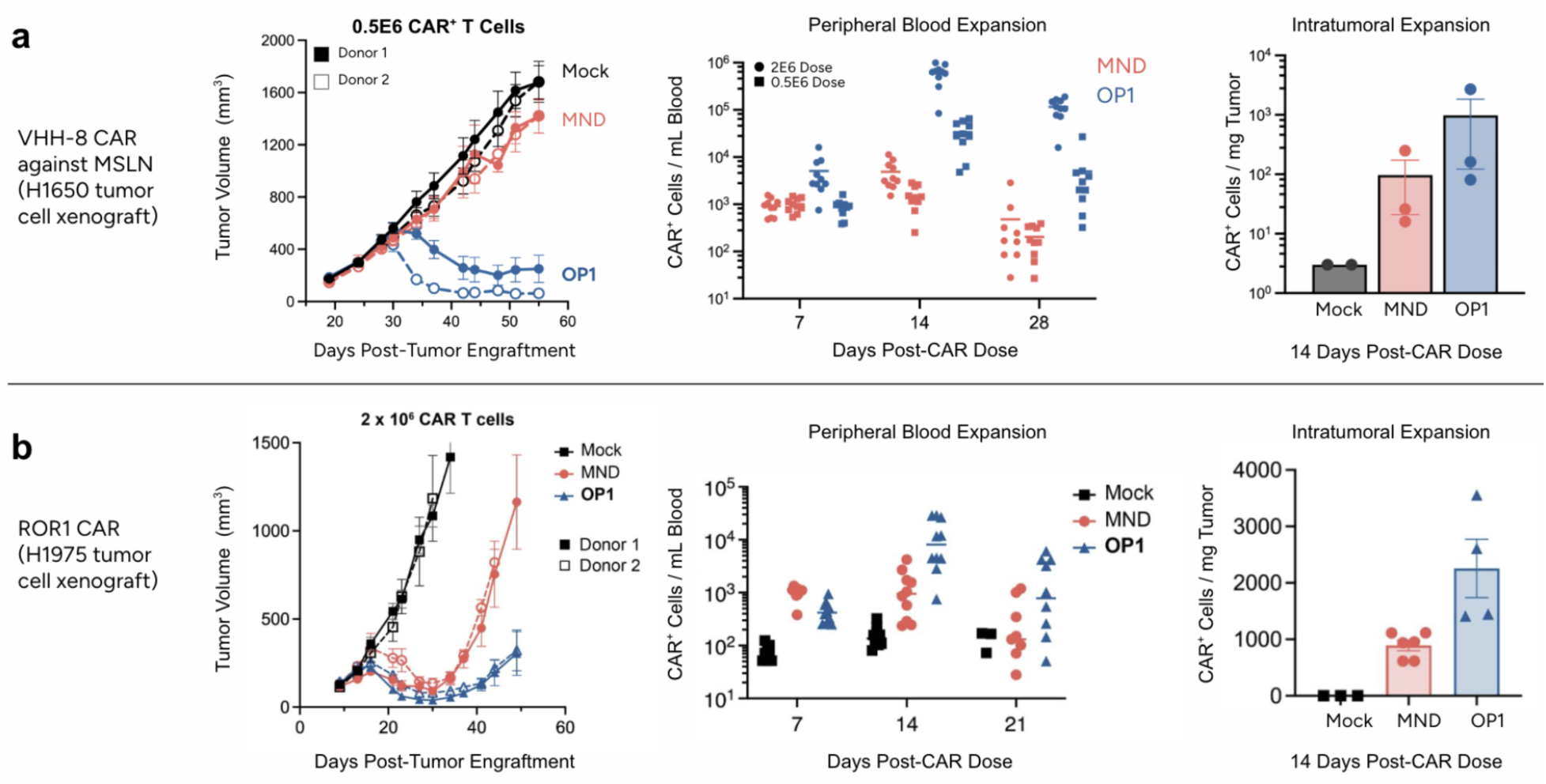
OP1-driven CAR T cells maintain better tumor control and CAR T cell expansion in both peripheral blood and intratumorally across two different solid tumor models. **a,** *in vivo* performance of OP1-driven VHH-8 MSLN CAR in H1650 xenografts. **b,** *in vivo* performance of OP1-driven ROR1 CAR in H1975 xenografts.

**Extended Data Figure 4.**
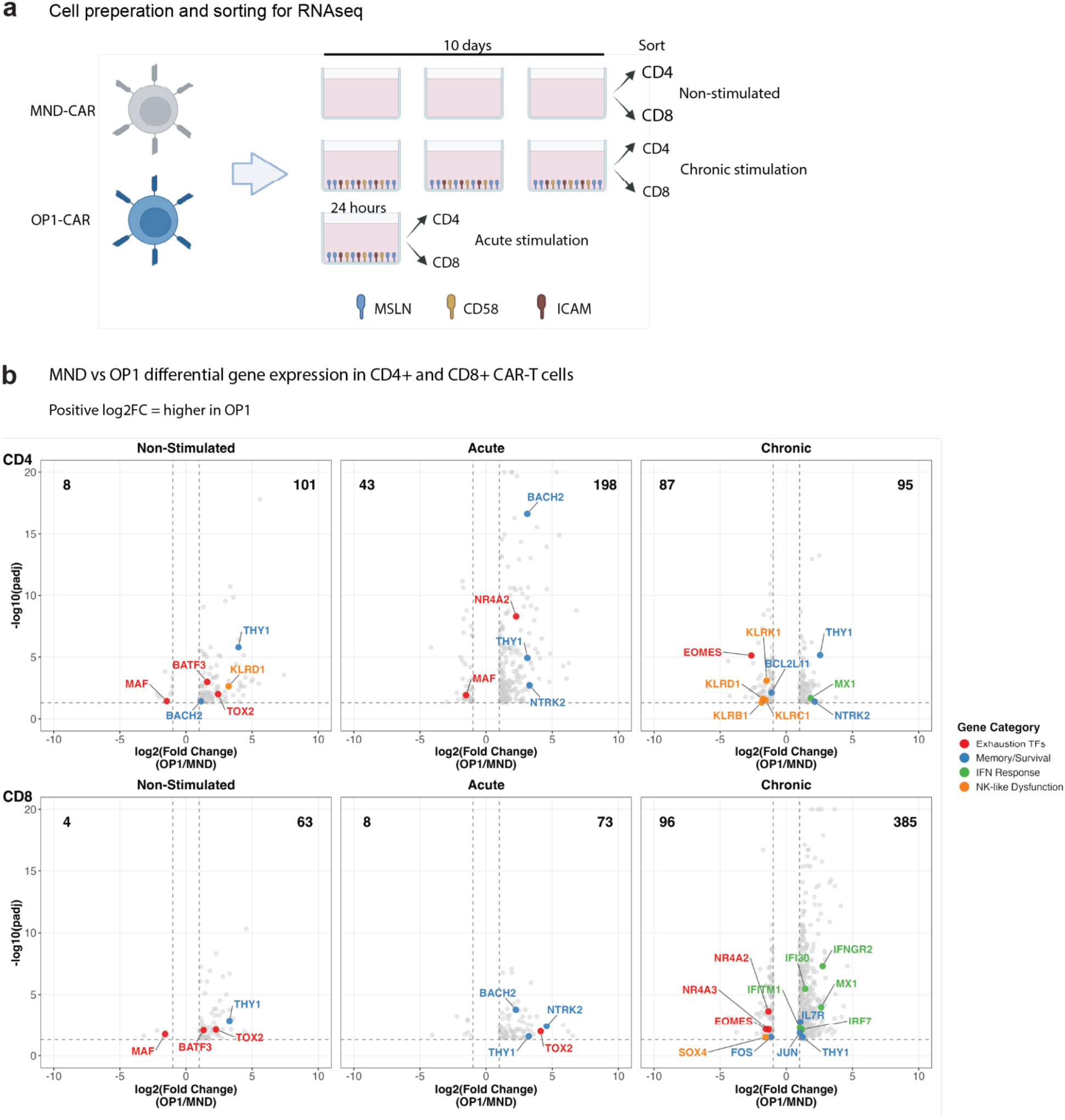
(goes with Figure 3). RNAseq analysis comparing OP1-driven versus MND-driven CAR T cells. **a**, schematic of CAR T cell preparation and sorting. **b**, data comparing differences in gene expression between OP1-driven and MND-driven CAR T cells (VHH-8 CAR) under non-stimulated, acute stimulation, or chronic stimulation conditions. Positive log2(fold-change) indicates genes higher in OP1N CAR; negative log2(fold-change) indicates genes higher in MND CAR. Illustrations in (a) were created using BioRender (https://biorender.com, Fontana, T. 2024).

**Extended Data Figure 5.**
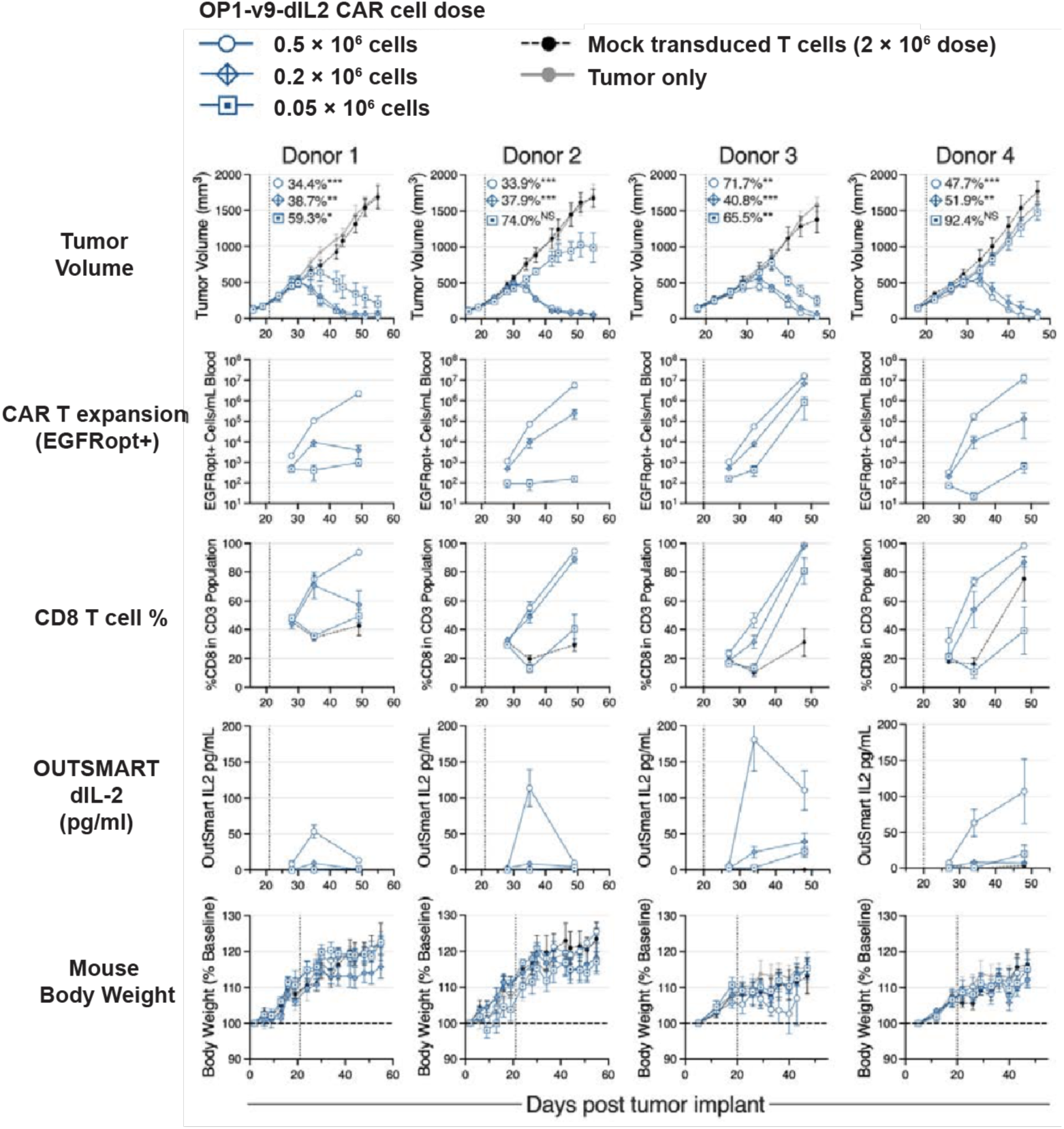
(goes with Figure 4c-d). **Data from additional mice and all 4 independent blood donors for subcutaneous H1650 xenograft studies**: tumor volume, CAR T peripheral blood expansion (EGFRopt+ cells), % CD8 T cells among the human CD3 T cell population, OUTSMART dIL-2 levels in peripheral blood, and mouse body weight days post subcutaneous NCI-H1650 implant in NSG mice (5 per group, mean ± standard error shown). CAR T generated with 4 individual healthy donors. Mice dosed with indicated number of mock transduced T cells or CAR T at day 20 or 21 post tumor implant (vertical dotted line). Tumor burden over study duration is summarized as a ratio of area under the curve (AUC) averaged across all mice in each group compared to the average AUC for the mock transduction group controlling for baseline tumor volume, where 100% represents equal tumor burden to the mock group, and lower percentages represent lower tumor burden. P-value significance for tumor burden in relation to mock transduced T cells. NS = not significant, * p < 0.05, **p < 5E-3, ***p < 5E-4.

**Extended Data Figure 6.**
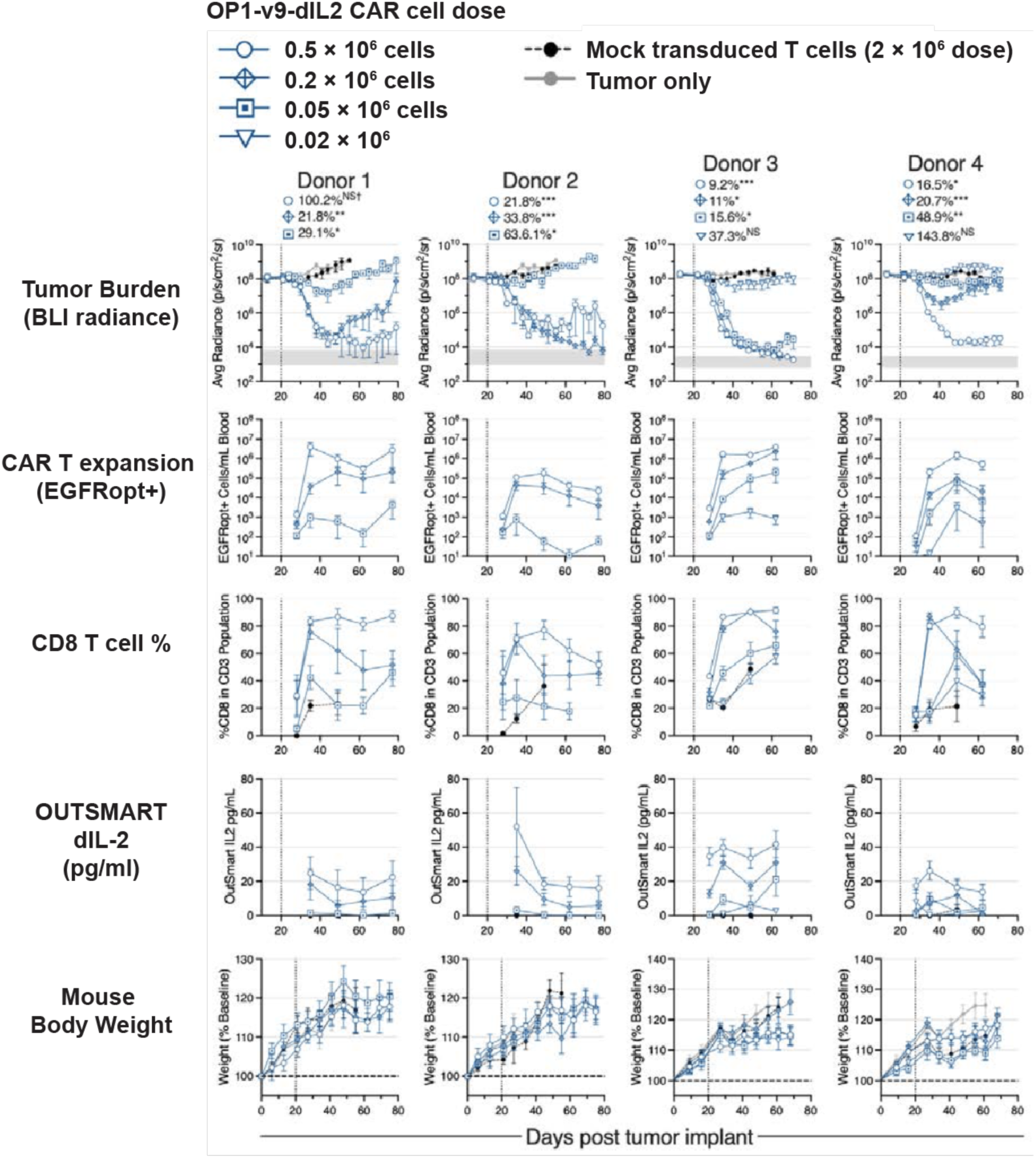
(goes with Figure 4e-f). **Data from additional mice and all 4 independent blood donors from intraperitoneal SKOV3 ovarian cancer models:** tumor burden (BLI radiance), CAR T peripheral blood expansion (EGFRopt+ cells), % CD8 T cells among the human CD3 T cell population, OUTSMART dIL-2 levels in peripheral blood, and mouse body weight days post subcutaneous SKOV3 implant in NSG mice (5 per group donors 1 and 2, 8 per group donors 3 and 4, mean ± standard error shown). CAR T generated with 4 individual healthy donors. Mice dosed with indicated number of mock transduced T cells or CAR T at day 20 post tumor implant (vertical dotted line). Tumor burden over study duration is summarized as a ratio of area under the curve (AUC) averaged across all mice in each group compared to the average AUC for the mock transduction group controlling for baseline tumor burden, where 100% represents equal tumor burden to the mock group, and lower percentages represent lower tumor burden. Gray area in tumor burden plots represents the background BLI levels from tumor naïve mice. P-value significance for tumor burden in relation to mock transduced T cells. NS = not significant, * p < 0.05, **p < 5E-3, ***p < 5E-4. † One animal was removed from study on day 38 post tumor implant, data is analyzed for statistical significance only up to this point for this group.

**Extended Data Fig. 7.**
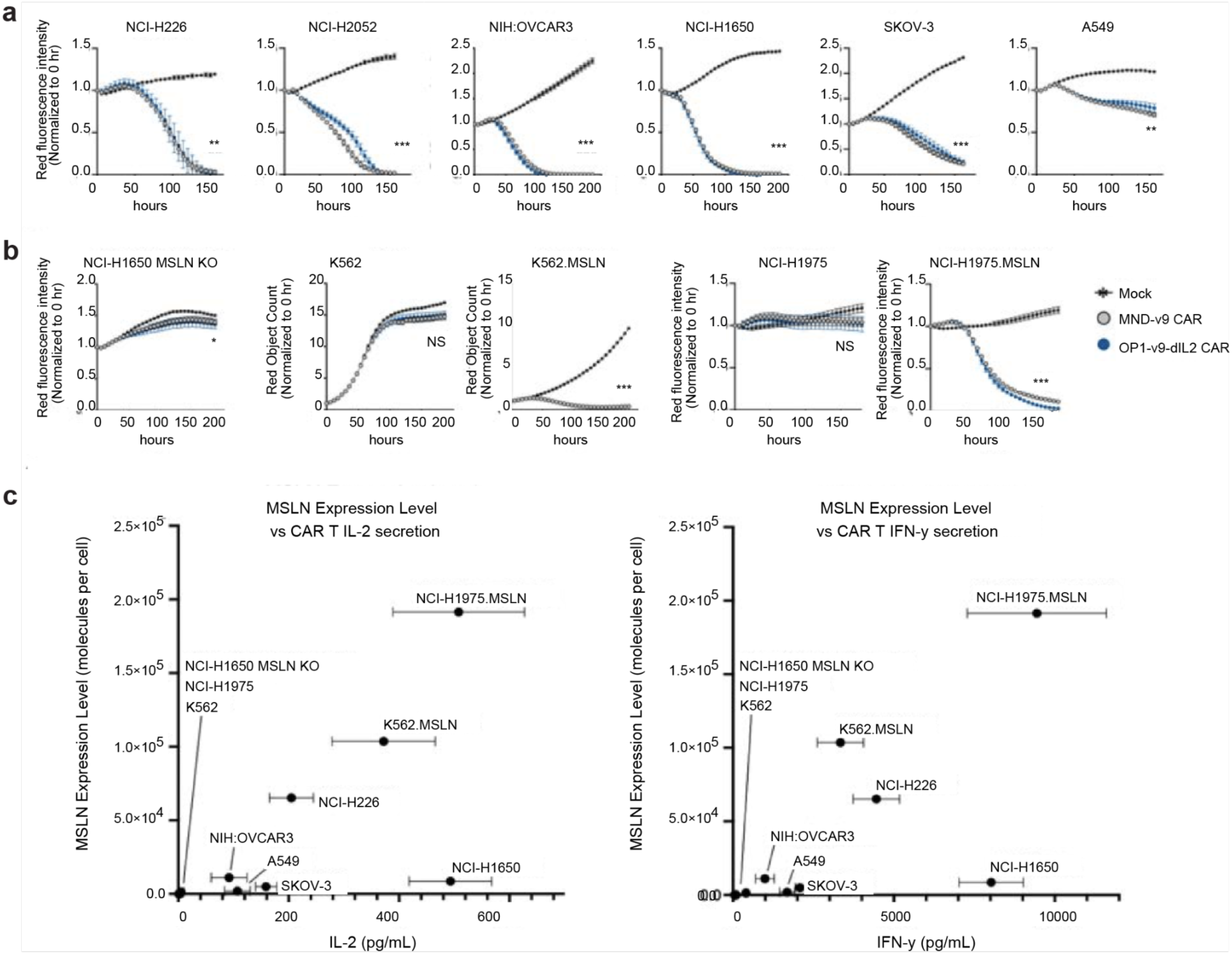
The CAR platform is highly specific and does not show cytotoxicity against low-MSLN-expression cell lines. MND-v9 CAR (gray) and full OP1-v9-dIL2 CAR construct (blue) show reduced killing of low-antigen cell lines: **a**, naturally-expressing MSLN expressing cell lines; **b,** engineered MSLN knockout (KO) or over-expression cell lines; measured by Incucyte live cell microscopy; p-value significance in relation to mock transduced T cells, NS = not significant, * p < 0.05, **p < 5E-3, ***p < 5E-4. **c,** Correlation of IL-2 or IFN-γ secretion by v9 CAR T cells in response to different MSLN expression levels on indicated cell lines; data is average of CAR T made from 4 healthy donors ± standard error.

**Extended Data Fig. 8.**
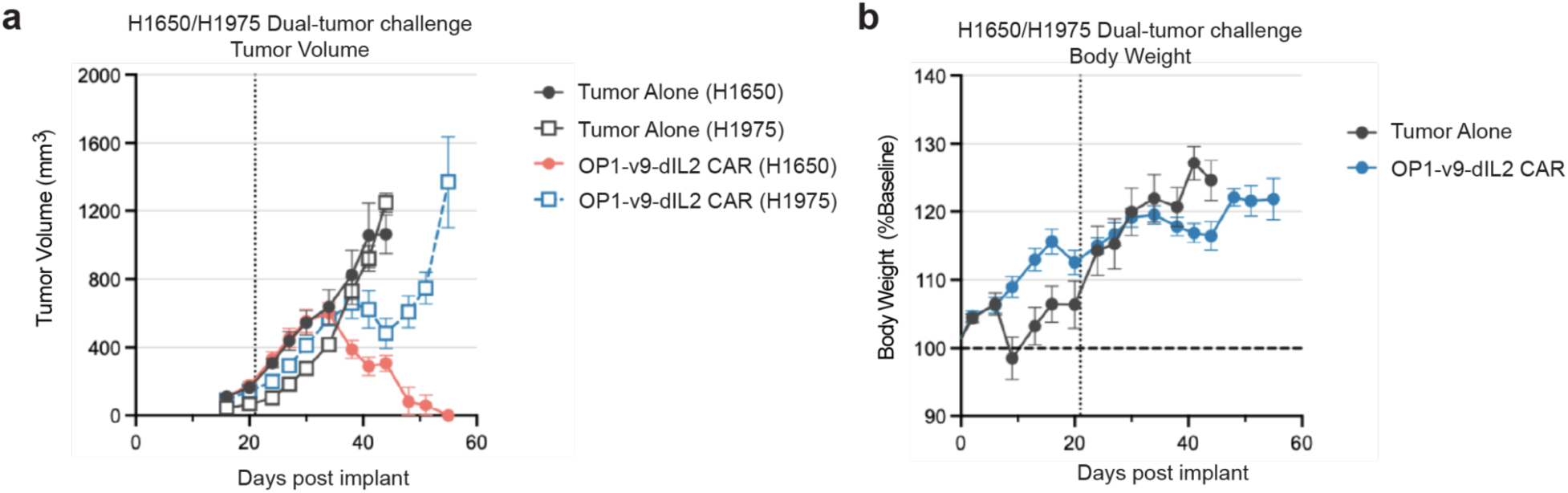
In a dual tumor model, OP1-v9-dIL2 CAR T cells specifically target MSLN+ H1650 tumors while sparing MSLN-H1975 tumors.

**Extended Data Figure 9.**
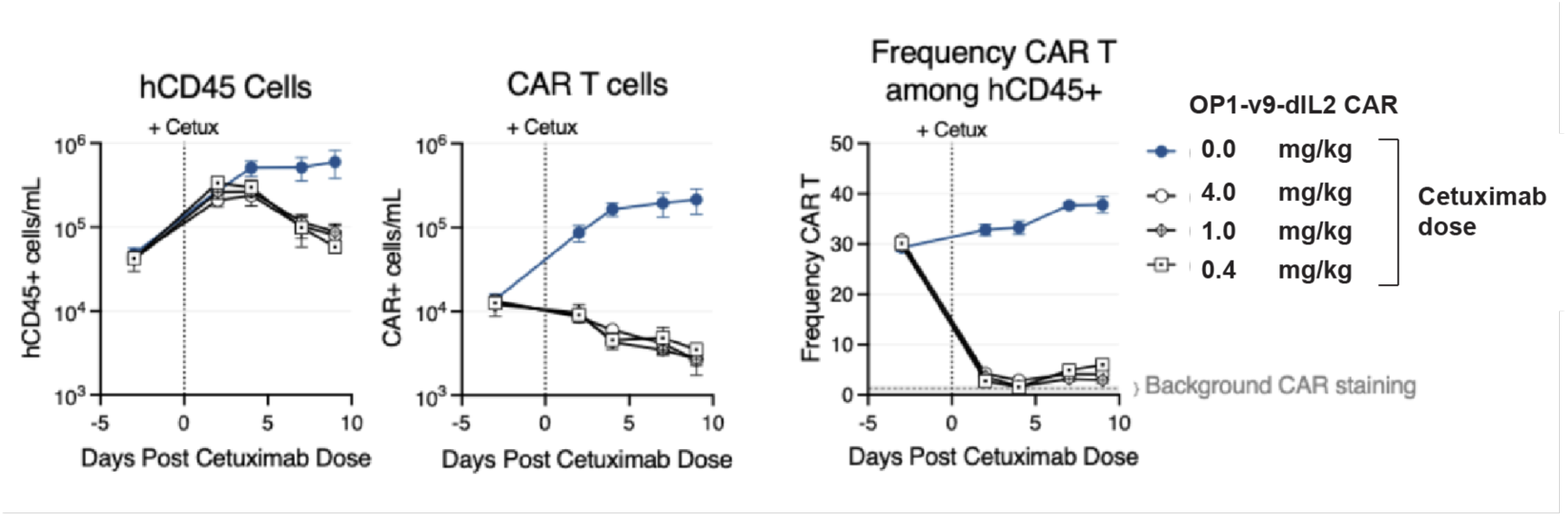
In vivo performance of the EGFRopt safety switch in response to Cetuximab (Cetux) in context of the full OP1-v9-dIL2 construct. Absolute counts of total human cells (CD45 positive) or CAR T cells (anti-G4S positive) in peripheral blood of NSG mice prior to and after dosing with the given concentration of Cetuximab. Frequency of CAR T cells among all human CD45 cells over the same time course; background level of CAR staining from G4S antibody indicated.

## MATERIALS AND METHODS

### Cell lines

NCI-H1650 (lung adenocarcinoma), SKOV3 (ovarian adenocarcinoma), NCI-H226 (lung mesothelioma), NIH:OVCAR3 (ovarian adenocarcinoma), K562 (chronic myeloid leukemia), H1975 (lung adenocarcinoma), and NCI-H2052 (lung mesothelioma) cell lines were obtained from ATCC. All cell lines were maintained according to ATCC recommendations and routinely tested for mycoplasma. Tumor cell lines for in vitro killing assays were engineered to express the red fluorescent protein mKate2. MSLN knockout and ectopic expression lines were generated from parental lines as described previously (see Supplementary Methods).

### VHH and scFv binder characterization

Parental muSS1 and humanized SS1 scFv-Fc protein test articles were expressed by Genscript on their TurboCHO platform, purified by ProA affinity and size-exclusion chromatography. Protein quality was confirmed by HPLC-SEC and SDS-PAGE. Thermal stability of WT and humanized SS1 proteins was evaluated using GloMelt dye on a Bio-Rad CFX384 Real-Time System instrument. The melt was performed with the first hold at 25°C for 30 seconds, then increasing temperature in increments of 0.5°C with a 20 sec hold, up to 95°C. Measurements were taken at every 0.5°C step. Binding of muSS1 and humanized SS1 scFv-Fc proteins to purified human MSLN Iso2[296-580] in 10x Kinetics Buffer (Sartorius) was evaluated using Biolayer Interferometry (BLI) on an Octet RED96 system. The scFv-Fc proteins were first captured on Protein A (ProA) biosensors with a loading threshold of 1 nm. After establishing a baseline, biosensors were dipped into wells containing MSLN analyte titrations (500 nM down to 2.1 nM in a 3-fold dilution series) to measure association for 180 sec and subsequently dipped into a running buffer to measure dissociation for 480 sec. Buffer-subtracted binding sensorgrams were fitted globally using a 1:1 Langmuir model to estimate rates of association (ka), dissociation (kd), and affinity (KD). Measurements were performed in triplicate.

### Protein production and purification

MSLN ECD expression (OPR1501 MSLN Iso2-Avi-His):

Expi293 cells were thawed and cultured in shake flasks at 37 °C with 8% CO2 for 3 passages prior to transfection. Plasmids encoding the protein of interest tagged with an appropriate N-terminal signal peptide (human IgK, for example) under the control of the CMV promoter were transiently transfected using the Expifectamine transfection reagent. One day post-transfection Expifectamine Enhancer 1 and 2 were added to cultures, and five days post-transfection supernatants were harvested by spin for 10 minutes at 4,000g at 4°C in centrifuge tubes and subsequently filtered with 0.2 um PES syringe or vacuum filters and held at 4°C until purification.

#### MSLN ECD Purification (OPR1501 MSLN Iso2-Avi-His)

All purification steps were performed at 2-8°C using an AKTA Pure 150 FPLC (Cytiva). As appropriate to supernatant volume, a 1 or 5 mL HisTrap Excel (Cytiva) was equilibrated with 4 column volumes of Buffer A: 1X PBS pH 7.4 with 500 mM NaCl (Teknova). Supernatants were loaded onto the column followed by washing with 20 column volumes of wash buffer, where the percentage of Buffer B: 1X PBS pH 7.4 with 500 mM NaCl and 500 mM Imidazole (Teknova) was in the range of 0-4% (Buffer A 100-96%) followed by elution with a linear gradient from wash buffer to 100% Buffer B over 10 column volumes followed by 10 column volumes of 100% Buffer B. Elution fractions showing UV absorbance at 280 nm were analyzed by Reducing SDS-PAGE and pooled prior to concentration by centrifugation in 3K MWCO filters (Cytiva) and loaded onto a Superdex 75 10/300GL or a Superdex 200 Increase Hi-Load 16/600 column with an isocratic elution of 1X PBS pH 7.2 (Thermo Fisher). Elutions from the Superdex column showing UV absorbance at 280 nm were run on reducing SDS-PAGE and fractions containing the protein of interest were pooled and concentrated by centrifugation in 3K MWCO nanosep, microsep, or macrosep devices (Cytiva) as appropriate and filtered with 0.22 um PES syringe or microcentrifuge filters (Cytiva/Corning/Agilent). Aliquots were flash frozen in liquid nitrogen and stored at -80°C until use for assays or further characterization. Concentration determination was done by A280 on the nanodrop UV spectrophotometer. Analytical Size Exclusion with MALS was performed by loading 25-50 ug of protein onto an equilibrated UP-SW2000 (Tosoh), UP-SW3000 (Tosoh), or a 1.7 um Acquity Premier Protein SEC 250Å column (Waters) on an Agilent 1260 Infinity II HPLC equipped with a Wyatt Dawn MALS detector and OptiLab differential refractive index detector with 1X PBS pH 7.2 (Thermo Fisher) as the mobile phase to assess native molecular weight and oligomeric state. Proteins were also analyzed for purity by reducing and non reducing SDS-PAGE.

### Lentiviral construct designs

T cells were engineered with a self-inactivating, third-generation HIV-1–derived lentiviral vector encoding a single polycistronic transgene. The “OP1-v9-dIL2” construct is driven by an NF-κB–responsive inducible promoter (OP1), and encodes three products separated by 2A self-cleaving peptides: (i) a mesothelin-targeted CAR (humanized SS1-derived hSS1v9 scFv; IgA2-derived spacer, CD28 transmembrane domain, and 4-1BB and CD3ζ intracellular signaling domains), (ii) EGFRopt, a truncated human EGFR serving as a transduction marker and cetuximab-inducible safety switch, and (iii) Outsmart dIL2, a CD8α-targeted engineered IL-2. Additional vectors were engineered to include an alternative promoter (i.e., MND), various anti-MSLN binders (i.e., VHHs and scFvs) and transgenes that lack OUTSMART dIL2.

### Lentiviral vector production

Lentiviral vectors were produced via polyethylenimine (PEI)-mediated transient transfection of suspension-adapted HEK293T cells grown in FreeStyle™ 293 Expression Medium. Three days post-transfection, the lentiviral supernatant was harvested, clarified by centrifugation at 300 x g for 10 minutes, and subsequently sterile-filtered through a 0.22 µm filter. For flask-scale production, the clarified supernatant was concentrated by ultracentrifugation at 10,000 x g for 4 hours at 4°C over a 10% sucrose cushion. The resulting viral pellet was resuspended in T cell media supplemented with 5% Lenti-X PEG concentrator. For smaller, 2 mL-scale preparations, the supernatant was incubated with Lenti-X PEG concentrator (3:1 v/v) for 2 hours at 4°C and centrifuged at 3,000 x g for 2 hours to pellet the vector, which was then resuspended in T cell media. All concentrated viral stocks were aliquoted and stored at -80°C. Viral titers were determined by transducing primary T cells. Vector copy number (VCN) was quantified using a QIAcuity Digital PCR system. The calculation involved amplifying the genomically integrated lentiviral Rev Response Element (RRE), correcting for plasmid carryover by amplifying the plasmid origin (Ori), and normalizing to the cell number by amplifying the human albumin gene. Transducing units per milliliter (TU/mL) were subsequently calculated based on the VCN. This method was adapted from Jiang et al. (2015).

### CAR T cell production

Primary human T cells were isolated from healthy donor leukopaks (StemCell) by negative selection. T cells were activated using the TransAct reagent (Miltenyi Biotec) and transduced with lentivirus. CAR T cells were expanded for 7 days in G-Rex plates (Wilson Wolf) in OpTmizer T Cell Expansion SFM supplemented with Immune Cell Serum Replacement (Thermo Fisher), GlutaMAX, and human IL-2, IL-7, and IL-15 (R&D Systems). Transduction efficiency was determined by flow cytometry for the EGFRopt marker.

### CAR T Cell Stimulation with Immobilized Proteins

To assess CAR T cell activation and exhaustion, in vitro stimulation assays were performed using immobilized recombinant proteins, adapted from previously described methods^31–33^. Tissue culture plates were coated with 5 µg/mL of recombinant human mesothelin (MSLN), 2 µg/mL of recombinant human CD58-Fc, and 2 µg/mL of recombinant human ICAM-1-Fc (R&D Systems) in PBS. The plates were incubated for 2 hours at 37°C or overnight at 4°C. Following incubation, the coating solution was removed. For acute stimulation, CAR T cells were seeded onto the coated plates at a density of 1-2 × 10⁶ cells/mL and incubated for 24 hours. For chronic stimulation, CAR T cells were cultured on the coated plates for a total of 9-10 days. To maintain continuous stimulation, cells were enumerated and replated at 1-2 × 10⁶ cells/mL onto freshly coated plates every 2-3 days, completing four rounds of stimulation. After both acute and chronic stimulation protocols, cells were harvested for downstream functional and phenotypic analysis.

### In vitro spheroid killing assay

H226 tumor cells expressing mKate2 were seeded in 96-well ultra-low attachment plates to form spheroids. CAR T cells were added at various effector-to-target ratios. Spheroid integrity was monitored by live-cell imaging (Incucyte, Sartorius), and killing was quantified by measuring the integrated red fluorescence intensity over time.

### Soluble MSLN and MUC16 assay in vitro assay

To assess the effect of soluble MSLN on CAR T cells, CAR T cells were co-cultured for 6 days with H1650 cells at a 1:4 ratio (2,500 CAR T cells:10,000 H1650 cells) in a 2D adherent format. At the time of setup, 10 µg/mL recombinant MSLN was spiked into culture. 24 Hours after co-culture setup, 20µL of supernatant was removed for MSD analysis of IFN-γ, IL-2, and TNF-α. To assess the effect of soluble MUC16 on CAR T cells, CAR T cells were co-cultured for 6 days with H2052 cells at a 1:4 ratio (2,500 CAR T cells:10,000 H2052 cells) in a 2D adherent format. At the time of setup, 1 µg/mL recombinant MUC16 was spiked into culture. 24 Hours after co-culture setup, 20µL of supernatant was removed for MSD analysis.

### In vitro exhaustion assay

For chronic stimulation, CAR T cells were co-cultured for 9-10 days on tissue culture plates coated with 5 µg/mL recombinant human MSLN and 2 µg/mL each of CD58-Fc and ICAM-1-Fc (R&D Systems) in OpTmizer T Cell Expansion SFM supplemented with Immune Cell Serum Replacement (Thermo Fisher), GlutaMAX, and human IL-2, IL-7, and IL-15 (R&D Systems). Cells were re-plated on freshly coated wells every 2-3 days.

### Flow cytometry

For in vitro analysis of cells by flow cytometry, cells were harvested and transferred to 96-well plates (50,000–400,000 cells/well), washed twice with Cell Staining Buffer (CSB, BioLegend), and incubated with antibody mixtures containing Live/Dead eFluor 780 viability dye and Brilliant Stain Buffer Plus for 30 min at 4°C in the dark. Cells were subsequently washed and fixed in Fluorofix (Biolegend) prior to acquisition on Ze5 (Bio-Rad), or Attune (ThermoFisher) flow cytometers. Compensation was performed using UltraComp eBeads, ViaComp beads, and single-antibody-stained controls, with fluorescence spillover matrices calculated either via Bio-Rad Everest automated compensation, Attune Cytometric Software, or applied post hoc in FlowJo. Antibody concentrations were optimized by titration, selecting dilutions that maximized stain index while maintaining high geometric mean fluorescence intensity (gMFI). All analysis was performed using FlowJo software.

For analysis of mouse peripheral blood, samples were collected into EDTA tubes (BD). For peripheral cytokine analysis, the plasma layer was isolated from whole blood by diluting 1:1 volume with PBS and centrifuging at 1000xG for 5 minutes. For flow cytometry, 50uL of whole blood was stained stained with antibodies against human lineage markers (CD45, CD3, CD4, CD8), memory and exhaustion markers (CD45RA, CD62L, PD-1, TIGIT, CD39), and CAR-specific markers (EGFR and recombinant MSLN protein). Precision counting beads (BioLegend) were used for absolute quantification. After staining, samples were RBC lysed and fixed, then acquired on a ZE5 flow cytometer (Bio-Rad) and analyzed using FlowJo software.

### RNA sequencing from CAR T cells

Rested or antigen-stimulated CAR T cells derived from five healthy donors were sorted on a SH800 cell sorter (Sony) for pure EGFRopt+CD4+ and EGFRopt+CD8+ CAR T cells. Sorted CAR-T cells were lysed in RLT buffer + 40 mM DTT (Qiagen RNeasy micro kit) and stored at -80°C. RNA was purified using RNeasy micro kits as per manufacturer’s instructions. Purified RNA was then sent externally to the company Novogene for next generation sequencing using their standard mRNA library preparation and sequencing protocol. Briefly, mRNA is enriched using poly A capture, the RNA is fragmented and then reverse transcribed into cDNA, and the library is prepared for paired end sequencing on an Illumina NovaSeq. RNA-seq data was obtained for 63 samples out of 70 samples. The remaining samples had insufficient RNA for library preparation. RNA-seq experiments used the VHH-8 MSLN CAR.

### Analysis of RNA-sequencing data

The resulting FASTQ sequences were processed using the nf-core/rnaseq pipeline ^34^ run with Nextflow version 23.04.3.5875. Briefly, adapters and low-quality sequences were removed from reads, along with genome contaminants and ribosomal RNA. Reads were aligned to the human genome assembly GRCh38 using STAR ^35^ and transcript abundances were quantified using Salmon ^36^. Salmon-estimated counts were rounded to integers for input into DESeq2. Differential gene expression analysis was performed using DESeq2 ^37^. Genes with fewer than 300 total counts across all samples were excluded prior to analysis. The DESeq2 model was specified as ∼condition, where condition represents each unique combination of cell type (CD4 or CD8), CAR construct (MND or OP1), and stimulation state (non-stimulated, acute, or chronic, as described above). Differentially expressed genes were identified using Wald tests with Benjamini-Hochberg correction for multiple comparisons, with significance thresholds of an adjusted p-value < 0.05 and an absolute log2 fold change > 1 (corresponding to a fold change > 2). Pairwise comparisons between MND and OP1 were performed separately for each cell type and stimulation condition, yielding six contrasts. Differentially expressed genes were annotated by functional category, including exhaustion transcription factors, memory and survival markers, interferon response genes, and NK-like dysfunction markers. Results were visualized as volcano plots with genes colored by their functional category.

### Animal studies

All animal studies were conducted under an IACUC-approved protocol. Female NSG mice (The Jackson Laboratory) were implanted subcutaneously on the right flank with 5 × 10^6^ NCI-H1650 cells mixed 1:1 with Matrigel (Corning). For the ovarian cancer model, 5 × 10^6^ SKOV3-ffLuc-GFP cells were implanted intraperitoneally and mixed 1:1 with Matrigel. When subcutaneous tumors reached approximately 150-200 mm^3, mice were randomized and treated with a single intravenous injection of CAR T cells. Tumor volume was measured twice weekly by digital calipers. For intraperitoneal tumors, CAR T cells were dosed 20 days after implant. Tumor burden was tracked using bioluminescent imaging by intraperitoneal injection of 150uL of 15mg/mL IVISbrite d-luciferin (Revvity) and images acquired using an IVIS Lumina LT (Revvity).

### In vivo evaluation of EGFRopt safety switch in response to Cexutimab

OP1-v9-dIL2 CAR T cells were administered to NSG mice and following engraftment, cetuximab was administered once intraperitoneally at doses of 4, 1 and 0.4 mg/kg. Though NSG mice lack T and B cells, the strain retains macrophages that can eliminate donor cells through ADCP and cetuximab contains a mouse IgG1 constant region capable of interacting with mouse Fcg receptors (Upton et al. 2021). Peripheral blood from treated animals (n = 5 per group) was assessed by flow cytometry for total engrafted human CD45 cells and CAR T cells (detected using an anti-CAR antibody, G4S).

### Statistical analysis

Statistical analyses were performed using R (v4.3.3) and GraphPad Prism (v10.2.2). For in vitro and in vivo killing assays, tumor burden was summarized by calculating the area under the curve (AUC). For studies with three or more donors, a linear mixed-effects model was used to compare treatment groups, with donors as a random effect. Survival curves were analyzed using the log-rank (Mantel-Cox) test. All other comparisons were made using two-tailed, paired Student’s t-tests on log10-transformed data. A p-value < 0.05 was considered significant.

#### Flow cytometry reagents used in this study

| Target | Fluorophore | Vendor | Catalog # | Clone |
| --- | --- | --- | --- | --- |
| **CD27** | BUV737 | BD Biosciences | 612829 | **L128** |
| **CD28** | BV711 | BioLegend | 302948 | **CD28.2** |
| **CD3** | BUV805 | BD Biosciences | 612893 | **UCHT1** |
| **CD3** | BV650 | BioLegend | 317324 | **OKT3** |
| **CD39** | BV510 | BioLegend | 328219 | **A1** |
| **CD4** | BUV496 | BD Biosciences | 612936 | **SK3** |
| **CD45** | PE-Dazzle 594 | BioLegend | 368530 | **2D1** |
| **CD45** | APC | BioLegend | 368512 | **2D1** |
| **CD45** | FITC | BioLegend | 368508 | **2D1** |
| **CD45RA** | BUV395 | BD Biosciences | 740298 | **HI100** |
| **CD56** | PE | BioLegend | 318306 | **HCD56** |
| **CD62L** | BUV615 | Thermo Fisher | 366-0629-41 | **DREG-56** |
| **CD8** | AF532 |  | 58-0088-42 | **RPA-T8** |
| **CD8** | BV786 | BD Biosciences | 563824 | **RPA-T8** |
| **CD8** | BV785 | BioLegend | 301046 | **RPA-T8** |
| **EGFR** | AF488 | BioLegend | 352908 | **AY13** |
| **EGFR** | BV711 | BioLegend | 352920 | **AY13** |
| **EGFR** | BV421 | BioLegend | 352911 | **AY13** |
| **LAG3** | SB780 | Thermo Fisher | 78-2239-42 | **3DS223H** |
| **PD-1** | PE-Cy7 | Thermo Fisher | 25-2799-42 | **eBioJ105 (J105)** |
| **TIGIT** | PE | BioLegend | 372704 | **A15153G** |
| **Tim-3** | BV421 | BioLegend | 364808 | **F38-2E2** |
| **MSLN**<br>**(Mesothelin)** | AF647 | Outpace | OFRE-016 |  |
| **Live-Dead-**<br>**eFluor 780** |  | Thermo Fisher | 65-0865-18 |  |

